# Title: AKI and CKD Worsen Outcomes in Experimental Sepsis: Creating a Reverse-translational Model of Acute Peritoneal Infection and Recovery in an Immunocompromised Rodent

**DOI:** 10.1101/2023.03.01.529424

**Authors:** Deana F. Floyd, James T. Colbert, Seth B. Furgeson, John R Montford

## Abstract

**Introduction:** Infection is a leading cause of morbidity and mortality in individuals with kidney disease. Both acute kidney injury (AKI), and chronic kidney disease (CKD) are clinical states that have been associated with higher risk of incident infection, and poor outcomes once infection has been established. A variety of host-and pathogen-specific factors are implicated as potential causes for these disparate outcomes including an altered host microbiome, innate and adaptive immune defects, and poor renal clearance and cytokines. However, there remains significant difficulty in modeling both human kidney disease and infection into an animal host. Likewise, there remains a poor understanding of the mechanisms underlying the unique immunodeficiency imparted by AKI and CKD, and if either condition imparts disparate risk.

**Methods:** C57BL/6J mice were given vehicle or aristolochic acid (AA) to create AKI (control, AKI groups) or CKD (control, CKD groups). Donor mice from all four groups underwent sterile cecal dissection and creation of cecal slurry (CS) preparations, which was later injected into separate mice in a matched host-recipient manner, at either high or lower doses. Animals were clinically monitored for either 24- or 72-hours after inoculation, then euthanized. Animal survival, sepsis severity, temperature, weights, and transcutaneous glomerular filtration rate (tGFR) were tracked longitudinally throughout the study. Histology for kidney injury, peripheral blood flow cytometry for leukocyte counts, plasma cytokines, and typical markers for organ injury were determined.

**Results:** Compared to controls, animals with AKI experienced much more severe sepsis across virtually all tracked metrics, and no animals with AKI survived high-dose CS injection past 24-hours. AKI mice manifested with a peripheral defect in leukocytes early after sepsis, with severe and persistent cytopenias, and a dramatically heightened early pro-inflammatory cytokine response. Septic CKD mice also had worse outcomes than controls, though less severe, and occurring later than in animals with AKI. Interestingly, animals with AKI had worse clinical outcomes and evidence of organ injury than mice with CKD at any dose or time-point after inoculation, despite a higher mean baseline measured GFR.

**Conclusions:** Rodents with established AKI and CKD experience worse clinical outcomes and organ injury versus controls in a CS model intraperitoneal live-bacterial infection. Additionally, mice with AKI experienced earlier and more severe morbidity and mortality than animals with CKD.

## Introduction

Infection is a leading cause of death and disability in individuals with chronic kidney disease (CKD). High rates of death and complications from pneumonia,^1^ sepsis,^2^ and more recently COVID-19 infection^3, 4^ have been appreciated in this at-risk population. In advanced CKD, the uremic state has long been recognized as immunomodulatory predisposing to acquired innate^5, 6^ and adaptive^7, 8^immune cellular dysfunction, and dysregulation of cytokine production.^9^ Separate from CKD, there is growing awareness that individuals with acute kidney injury (AKI) are also predisposed to incident infection, and very poor outcomes once infection has been established.^10–13^ AKI is a systemic hyperinflammatory state, and patients at the highest risk of subsequent death with AKI are known to have drastically upregulated plasma cytokine levels.^14^ Additionally, a variety of preclinical and clinical studies document specific cellular defects in mononuclear cells^15^, neutrophils^16, 17^, and T-cells.^18^ While there is growing recognition of the burden that infection poses to patients with AKI and CKD, there is little consensus on the basic mechanisms driving higher infection rates. Furthermore, there is few data supporting if AKI and CKD impart disparate risk; and if so, what are the drivers?

Preclinical modeling of AKI and CKD is difficult, and methods of creating kidney injury in a rodent have traditionally been used to study physiology and intrarenal injury/repair. Modeling infection in a rodent is also difficult, as injection of sterile irritants (such as lipopolysaccharide (LPS)) are known to poorly mimic human infection.^19^ Other methods, such as surgical cecal ligation and puncture (CLP, widely considered the “gold standard” of acute bacterial infection modeling) are highly variable, morbid, require surgery and anesthesia, and thus may be poorly suited for the study of a previously immunocompromised host. Recently, many investigators have utilized the cecal slurry (CS) infection model as an improved method to recreate acute bacterial peritonitis. CS peritonitis involves the inoculation of a recipient animal with homogenized donor stool derived from pooled separate hosts. The CS peritonitis method is reproducible, creates dose dependent sepsis severity, avoids the need for anesthesia and laparotomy, but is nonetheless a live bacterial polymicrobial model of infection.^20–22^

We aimed to establish a reproducible model of bacterial infection using CS injection in rodents with preexisting AKI and CKD to evaluate if existing kidney disease predisposes to worse sepsis outcomes and an altered early immune response to infection. We first established parallel rodent model of AKI and CKD using low dose aristolochic acid (AA), a phytotoxin causing human kidney disease in the developing world.^23, 24^ We then collected cecal contents from donor mice in these groups (along with paired control animals), created a glycerol-stabilized homogenous cecal slurry, and injected AKI, CKD, and control mice with the solution to create a model of acute bacterial peritonitis. *Our overall hypothesis was that mice with AKI and CKD would have worse clinical sepsis outcomes than control mice, having an altered early innate immune-mediated response to infection. Furthermore, we predicted that mice with preexisting CKD, having a much lower baseline GFR in our model than mice with AKI, would experience the worst clinical outcome after CS inoculation*.

## Results

### Establishing separate rodent models of AKI and CKD using low dose aristolochic acid administration

*AKI Model* C57BL6/J male mice were given either vehicle (N=10) or AA (N=10) at a dose of 2mg/kg intraperitoneal (i.p.) three times weekly for three weeks to create the model of indolent AKI (**Figure 1A**). Compared to vehicle-injected controls, mice with AKI experienced a gradual, but significant decline in measured transcutaneous glomerular filtration rate (tGFR) (**Figure 1B**) starting at 1 week after initial injection of AA; culminating with a mean reduction in tGFR by nearly half of controls by day 21 (537 vs 966 μL/min/100g BW, p<0.0001, AKI vs controls, respectively). AKI animals had also lost significant weight versus controls (24.6 vs 29.0 g body weight, p=0.04), experienced slight anemia (13.6 vs 15.6 g/dL hemoglobin, p<0.0001), and had evidence of nephrosarca (0.75 vs 0.62 % wet kidney weight/body weight, p<0.0001). Plasma BUN was unchanged, though creatinine was significantly elevated in the AKI mice vs controls (0.35 vs 0.14 mg/dL, p<0.0001). Histologic examination of kidneys from AKI mice showed preservation of glomeruli with moderate acute tubular injury (ATI); characterized by dilated and simplified tubular lumens, an expanded, edematous cortex with evidence of mild acute interstitial nephritis, and a slight loss of volume in the outer medullary stripe (3.3 vs 0.1 acute tubular injury (ATI) score, p<0.0001, AKI vs controls, respectively) (**Figure 1D**). Interstitial fibrosis was significantly increased in AKI versus control kidneys, although the magnitude of fibrosis was low (2.2 vs 0.7 % cortical fibrosis, p=0.002).

**Figure 1.**
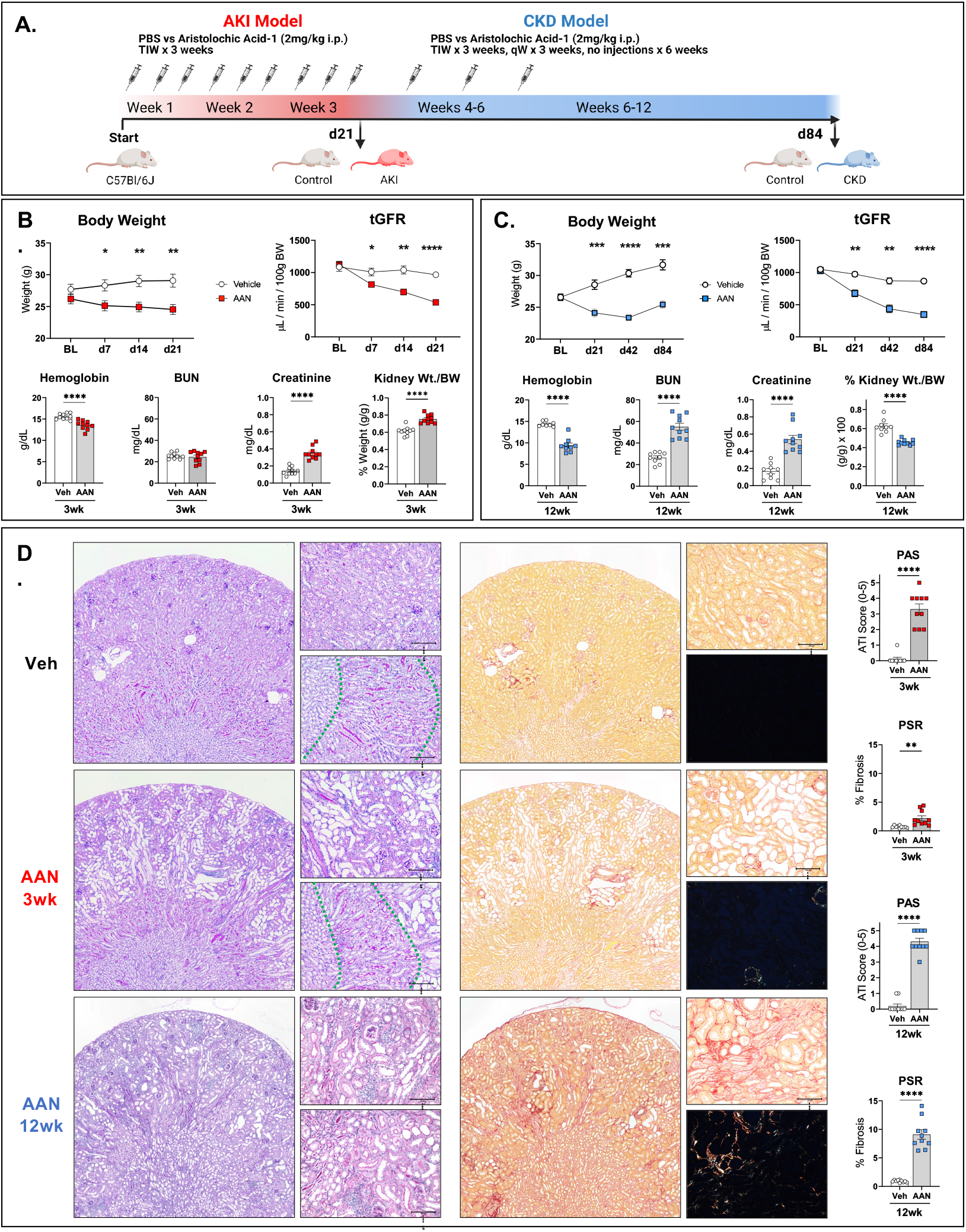
Development of Separate AKI and CKD models using Low Dose Aristolochic Acid Repeated Injections. A) Mice were injected with repeated vehicle (PBS) or aristolochic acid (AA) intraperitoneal injections and euthanized at either day 21 for AKI studies (N=10 vehicle, N=10 AAN) or day 84 for CKD studies(N=9 vehicle, N=10 AAN). In AKI (B) and CKD studies (C) serial body weight, tGFR trends are shown, along with whole blood hemoglobin, plasma BUN, Creatinine, and kidney weight/body weight at terminal collection. D) Histologic examination of kidney tissues from a representative vehicle control mouse and AKI and CKD mouse are shown under 2x and 20x magnification. PAS stained tissues (left panels) and PSR stained tissues with corresponding polarized 20x image (right panels) are shown. Quantification of PAS and PSR stained tissues from affected animals were determined (right most graphs). * p<0.05, **p<0.01, ***p<0.001, ****p<0.0001.

*CKD Model* Separate groups of vehicle (N=9) or AA-injected (N=10) mice were given a similar initial AKI protocol, but then underwent isolated weekly AA injections for 3 more weeks, allowed to recover for an additional 6 weeks, and were euthanized on day 84 (12 weeks after the first injection) (**Figure 1A**). CKD mice experienced a similar indolent AKI phase as in the AKI model, with similar weight changes and loss of tGFR during the first 21 days of injury (**Figure 1C**). By the end of the study, CKD mice regained some weight but remained significantly smaller than age-matched controls (25.4 vs 31.6 g body weight, p=0.0007, CKD vs controls, respectively). However, there were notable clinical, biochemical, and kidney histologic differences in CKD-vs AKI-mice. tGFR continued to significantly decline in CKD mice over time, to a mean of 348 vs 847 μL/min/100g BW in control mice (p<0.0001) by day 84. By 12 weeks, plasma BUN (55.0 vs 26.4 mg/dL, p<0.0001) and creatinine (0.54 vs 0.17 mg/dL, p<0.0001) were significantly elevated versus controls and hemoglobin concentration was significantly depressed versus controls (9.4 vs 14.4 g/dL, p<0.0001). Kidney atrophy, not edema, was appreciated in kidney weights of CKD mice (0.63 vs 0.46 %kidney weight/body weight, p<0.0001). Histologic sections again revealed preserved glomeruli, but unlike kidneys from AKI mice, CKD mouse kidneys had profound tubular atrophy, nephron dropout, widespread tubulointerstital nephritis, proteinaceous cast formation, and “medullification” of entire cortical regions (**Figure 1D**). Also, unlike kidneys from AKI mice, CKD kidney specimens showed significantly increased interstitial fibrosis vs controls (9.1 vs 0.9% cortical fibrosis, p<0.000).

### Mice with pre-existing AKI experience heightened sepsis severity versus controls using a novel method of cecal slurry peritonitis

**Figure 2** outlines the experimental approach used in CS sepsis studies. Vehicle control (N=30, hereafter “No-AKI” controls) and AA-injected AKI mice (N=30) were generated as before. Baseline (after 21 days of vehicle or AA-injections, and immediately before sepsis studies commenced) weights (**Figure 3A**) and tGFRs (**Figure 3B**) were similar to animals described in original model development. A sham solution (PBS/10% glycerol, 0.3mL), low dose cecal slurry (CS-LD, 0.2mL or 20mg) or high dose cecal slurry (CS-HD, 0.3mL or 30mg) was administered in a matched donor/recipient group manner (No-AKI CS into No-AKI mice, AKI CS into AKI mice) as a single i.p. injection and mice monitored clinically with a validated clinical sepsis score, rectal temperature measurements, and daily weights for either 24- or 72-hours. No-AKI controls, the sham, CS-LD, and CS-HD animals all had 100% survival until their designated endpoint collections. However, animals in the AKI group experienced high attrition. One animal in the AKI group receiving CS-LD was euthanized for moribund status (t=24h after CS inoculation), and all animals administered CS-HD did not survive past 24 hours (three animals with non-survival by 20h; and remaining euthanized at 20-24h for moribund status, **Figure 3C**). All animals had a graded response to CS doses as measured by clinical sepsis scores (**Figure 3D**) and degree of hypothermia (**Figure 3E**). However, compared to No-AKI controls, animals with AKI had significantly worse sepsis severity and hypothermia by as early as 4 hours after inoculation, even when receiving the lower CS dose (8.7 vs 5.5 4- hour sepsis score, p=0.0038, AKI vs control mice, respectively).

**Figure 2.**
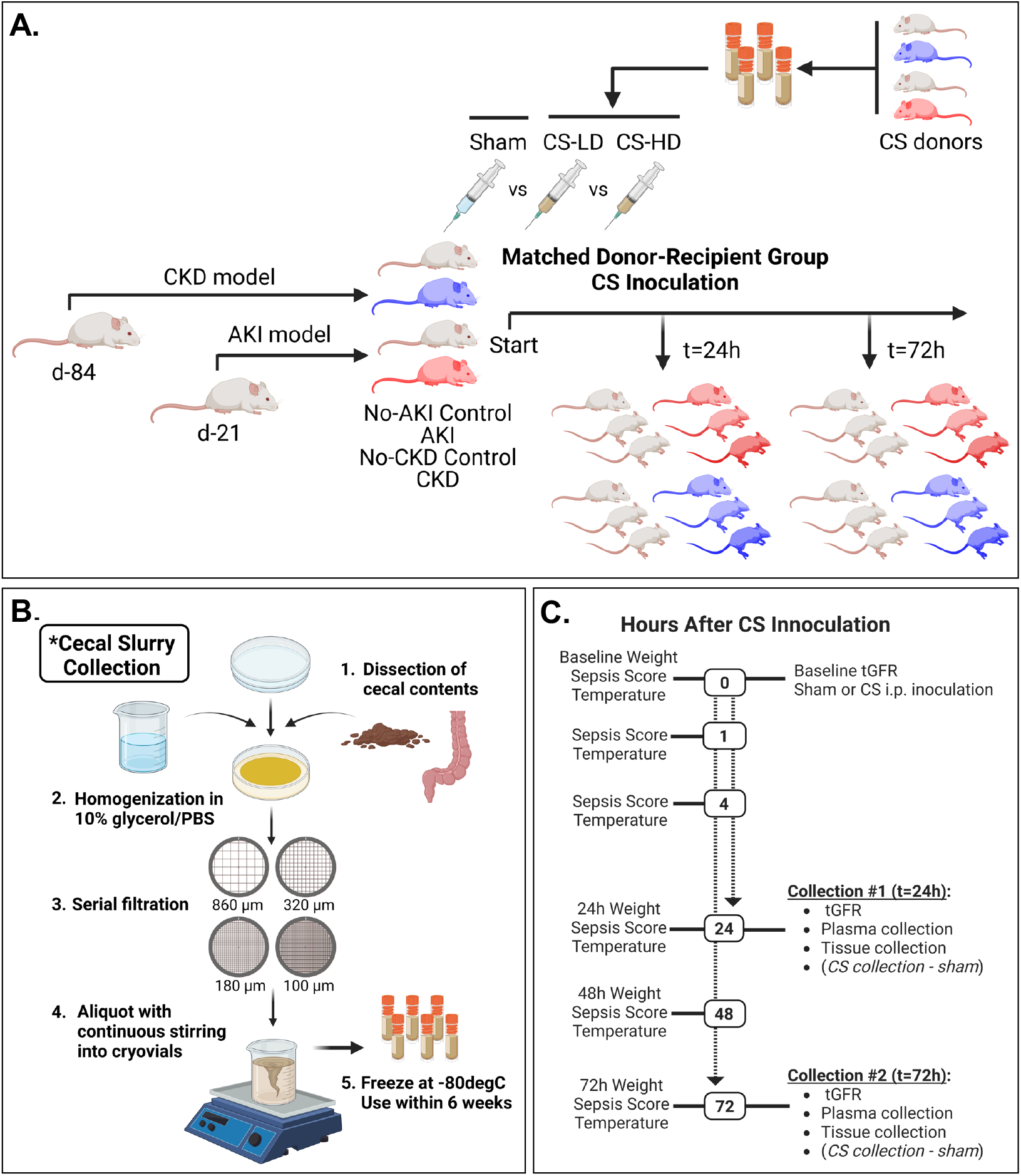
Design of CS Studies. A) Outline of study design. AKI or CKD groups generated, animals receiving sham injections also served as CS donors. B) Creation of CS host-matched stock solutions. (C) Diagram of post-CS inoculation clinical monitoring and endpoint analyses.

**Figure 3.**
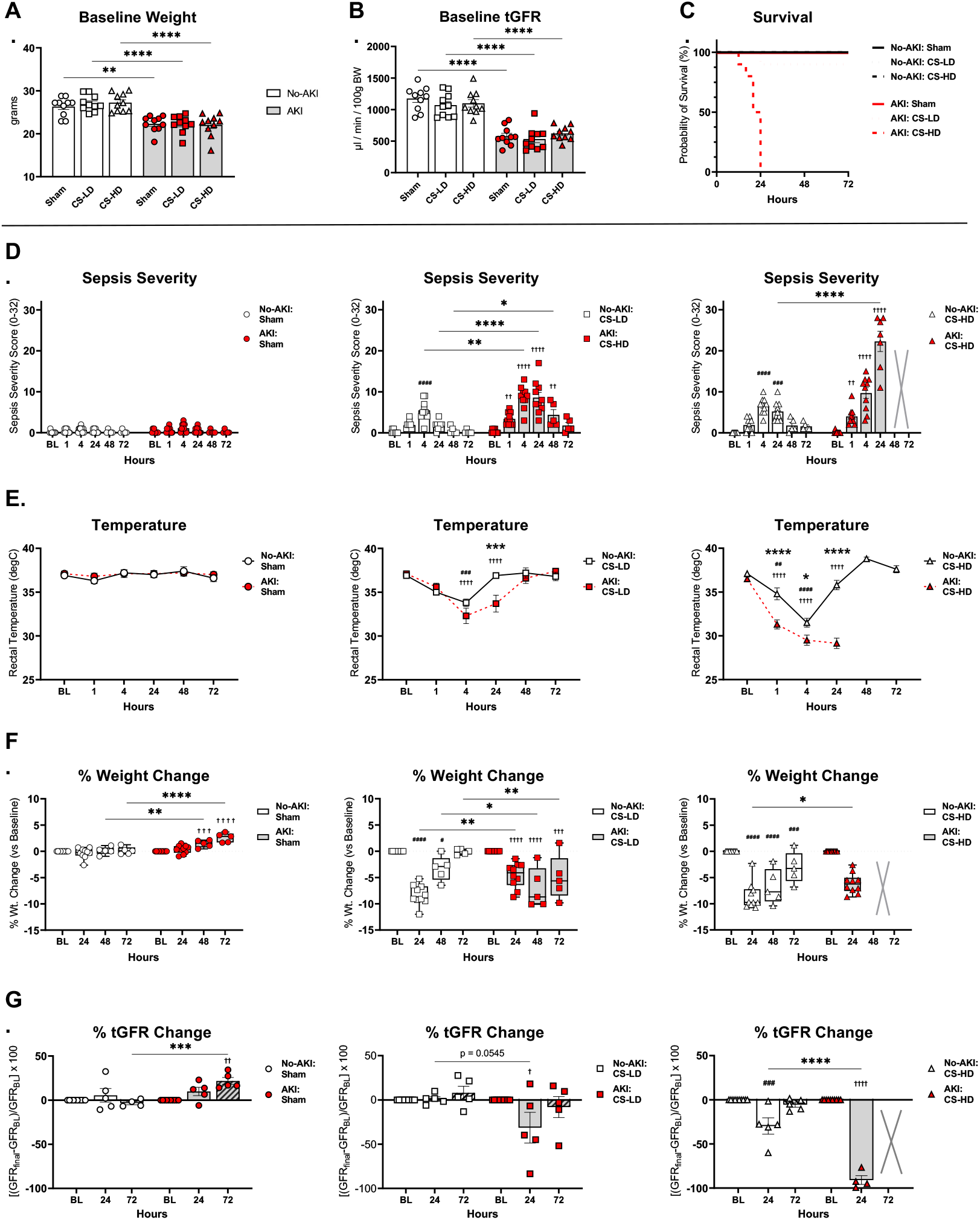
AKI CS Sepsis Studies. A) Baseline weights and B) tGFRs are shown for No-AKI (N=30) and AKI (N=30) mice. C) AKI animals suffered early, severe mortality in high dose groups, No-AKI animals had 100% survival in all groups. D) Serial assessments of clinical sepsis severity, E) rectal temperature, F) change in weight after inoculation, and G) change in tGFR was assessed in No-AKI and AKI mice undergoing sham injections (D-G, left panels), CS-LD injections (D-G, middle panels), and CS-HD injections (D-G, right panels). Compared to No-AKI controls, AKI mice had early worsening of sepsis severity and hypothermia. AKI mice also had lower kidney function 24h after low-and high-doses of CS. No-AKI mice lost the most weight at 24h, but steadily regained weight throughout the study. (Within No-AKI group: # p<0.05, ##p<0.01, ###p<0.001, ####p<0.0001; Within AKI group: †p<0.05, ††p<0.01, †††p<0.001, ††††p<0.0001; No-AKI vs AKI group: * p<0.05, **p<0.01, ***p<0.001, ****p<0.0001)

Mice without AKI demonstrated a reduction in 5-10% of baseline body weight, peaking at 24 hours after CS inoculation, with most regaining body weight by 72 hours (**Figure 3F**). Interestingly, in mice with AKI there was significantly less acute weight loss than in No-AKI mice at 24hrs after LD-CS (−4.6 vs −8.3% weight loss, p<0.007) and a similar trend after HD-CS (−6.1 vs −8.5% weight loss, p=0.09). However, surviving mice with AKI experienced a comparative inability ability to regain body weight by 72 hours after LD-CS (−5.0 vs −0.8% weight loss, p=0.008). Only mice with AKI given sham-injections gained weight throughout the study, most likely resulting from the cessation of AA-injections and expected AKI recovery. Additionally, all recovered CS-HD AKI mice, and the one euthanized CS-LD AKI mouse had significant burdens of bacteremia (**Supplemental Figure 1**).

tGFRs were assessed in each animal at two time points: at baseline and immediately before planned terminal collection to determine if there was evidence of sepsis-induced new AKI in No-AKI mice, or “AKI-on-AKI” among mice with preexisting AKI (**Figures 3G, Supplemental Figure 2A**). As previously mentioned, No-AKI and AKI mice had significantly divergent tGFRs at baseline before sham or CS inoculations (**Figure 3B**). In No-AKI mice, kidney function was preserved after CS-LD injections, with only animals receiving CS-HD showing a non-significant trend toward reduced renal function 24-hours after inoculation vs baseline (747 vs 1044 µL/min/100g BW, p=0.11, respectively); which showed normalization vs baseline in the 72-hour group (1095 vs 1155 µL/min/100g BW, p=0.28, respectively). In mice with existing AKI there was a stronger, but non-significant, trend towards additive AKI at lower CS doses (372 vs 586 µL/min/100g BW, p=0.34 24-hours vs baseline). After CS-HD, most animals with preexisting AKI had little detectable kidney function (58 vs 564 µL/min/100g BW,p=0.0008, 24-hours vs baseline, respectively) appreciated by a “flat” tGFR curve (**Supplemental Figure 2C**). Of note, not all AKI mice receiving CS-HD underwent tGFR monitoring given the degree of sepsis severity (four out of the seven surviving mice by 20-24h, the remaining three immediately euthanized to minimize further distress).

### Mice with AKI experience more biochemical evidence of severe sepsis and organ injury versus No-AKI control mice receiving both low-and higher CS doses

Plasma BUN and creatinine largely mirrored the tGFR findings in No-AKI and AKI mice, with BUN showing the better early sensitivity reflecting functional renal decline in septic mice with preexisting AKI (**Figure 4A**). Compared with sham-injected No-AKI animals after 24-hours, only the No-AKI CS-HD groups had significantly increased BUN (64 vs 23 mg/dL, p=0.0001, respectively), but no changes in creatinine consistent with measured GFR. Animals with AKI showed more severe azotemia at both low-and high-CS doses having significantly increased BUN (55 and 101 vs 24 mg/dL, p=0.002 and <0.0001, respectively). Creatinine showed a delayed increase in AKI animals vs tGFR changes, peaking at 72h in the low-dose CS group, but was significantly elevated acutely in the surviving high CS dose groups at 20-24 hours.

**Figure 4.**
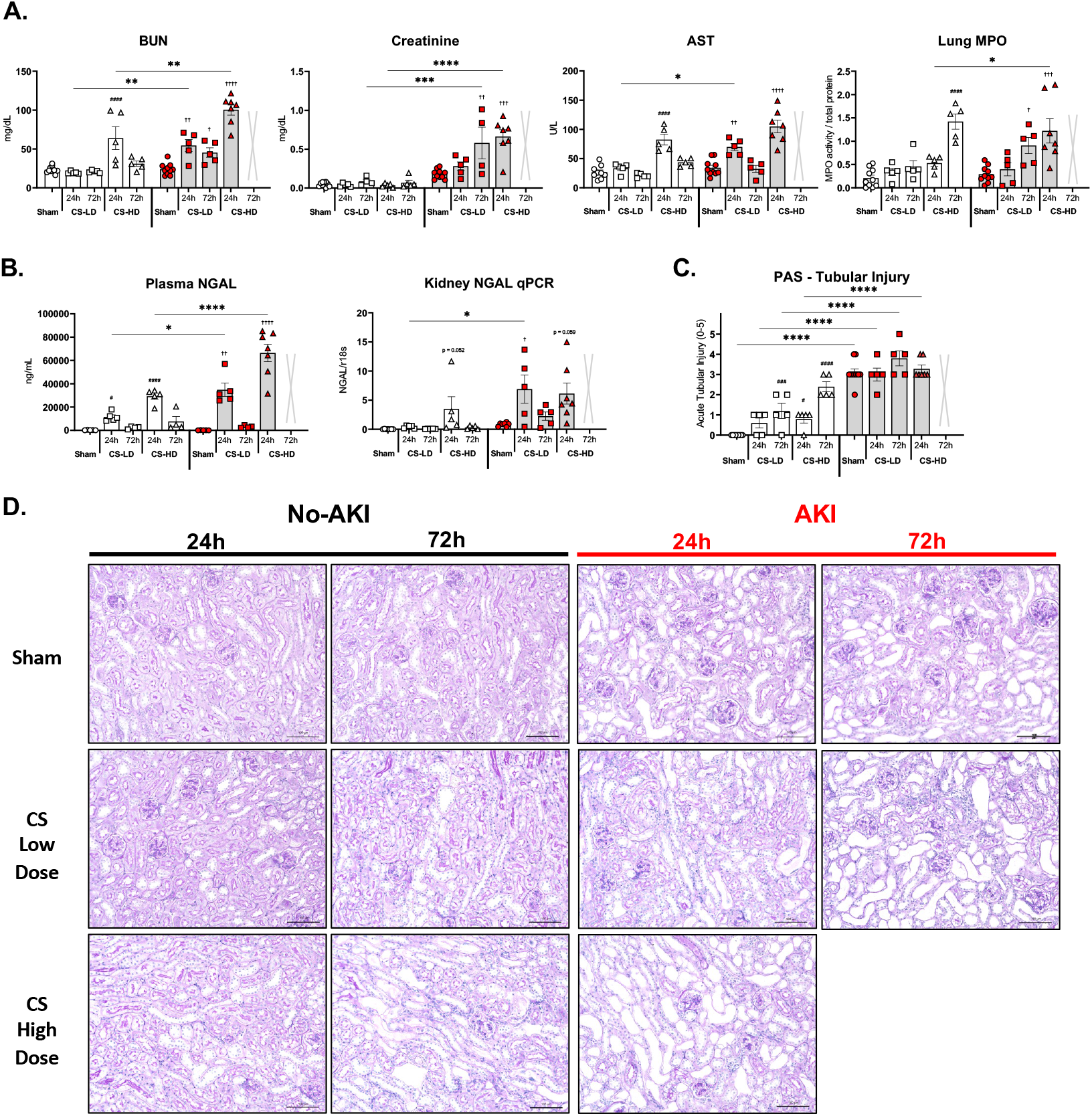
AKI mice show evidence of early and late organ dysfunction during sepsis. A) Plasma BUN, Creatinine, AST, and lung MPO levels were obtained revealing more azotemia and sepsis severity in AKI mice receiving either low or higher-doses of CS vs controls. B) NGAL levels were assessed using an ELISA in the plasma and via qPCR in the kidney, demonstrating significantly higher levels across the study in AKI mice. D) PAS images (20x) from cortical regions of affected animals demonstrate a delayed increase in acute tubular injury, peaking at 72 hours after CS inoculation. C) Quantification of ATI in mice with preexisting AKI was difficult to discern vs in No-AKI control mice. (Within No-AKI group: # p<0.05, ##p<0.01, ###p<0.001, ####p<0.0001; Within AKI group: †p<0.05, ††p<0.01, †††p<0.001, ††††p<0.0001; No-AKI vs AKI group: * p<0.05, **p<0.01, ***p<0.001, ****p<0.0001)

We also compared global sepsis biomarkers such as AST (**Figure 4A**) and plasma NGAL (**Figure 4B**) between the two groups. AST results paralleled those of BUN, and plasma NGAL performed more accurately as a sepsis biomarker, being significantly elevated with both low-and high-CS doses vs shams in the No-AKI group. Even so, mice with AKI had significantly higher plasma NGAL than control mice across the study at 24-hours (34852 vs 11635 ng/mL, p=0.02, CS-LD); 66466 vs 29325 ng/mL, p<0.0001, CS-HD, respectively). Plasma NGAL in AKI might come from many sources, including both the kidney and liver in an IL-6 dependent manner as we and others have previously published^25^. Therefore, we analyzed frozen kidney tissue for NGAL message by qPCR which showed a similar pattern of: upregulation only in CS-HD No-AKI animals, and significantly higher upregulation in both low-and higher-dosed AKI animals. Finally, frozen lung tissues were analyzed for MPO-enzymatic activity to determine if any remote lung injury from sepsis was present. Interestingly, lung MPO activity appeared to increase the most after 72-hours post-inoculation in a delayed fashion in recovering animals.

PAS-stained kidney sections from AKI mice and controls showed subtle changes in tubular injury assessed by a combination score of tubular simplification/dilation, presence of proteinaceous casts, and degree of interstitial nephritis (**Figure 4D**). Control mice showed significant increases in tubular simplification and slight interstitial nephritis peaking at 72-hours (ATI scores 1.2 and 2.4 vs 0, p=0.001 and p<0.0001, low-and high-dose sepsis vs sham, respectively) though overall degree of histologic injury was minor. Determining differences in histologic injury in mice with preexisting AKI and sepsis was much more difficult; and while higher-dose CS injection trended towards worse tubular injury, driven by objectively more dilated tubules and interstitial nephritis (**Figure 4C**), these results were not statistically different versus baseline degree of tubular injury in sham-injected animals.

### AKI predisposes to severe peripheral immune defects and an aggravated acute immune response after cecal slurry inoculation

Flow cytometry was performed on peripheral blood specimens from affected animals. CS injection induced a dose-and time-dependent leukopenia in No-AKI mice, and this was worsened in mice with AKI (**Figure 5A**). Specifically, populations of lymphocytes, granulocytes, and monocytes all acutely dropped then partially rebounded by 72 hours after CS injection in No-AKI mice. Among mice with AKI no, or very little, peripheral reconstitution was appreciated among surviving animals. In lymphoid cells in AKI mice (**Figure 5A, middle panel**), CD19+ B-cells acutely fell 24-hours after CS injection and failed to rebound by 72 hours in low dose mice. However, CD4+, CD8+, and γΔ T-cells all increased as a percentage of leukocytes in AKI mice vs non-AKI mice.

**Figure 5.**
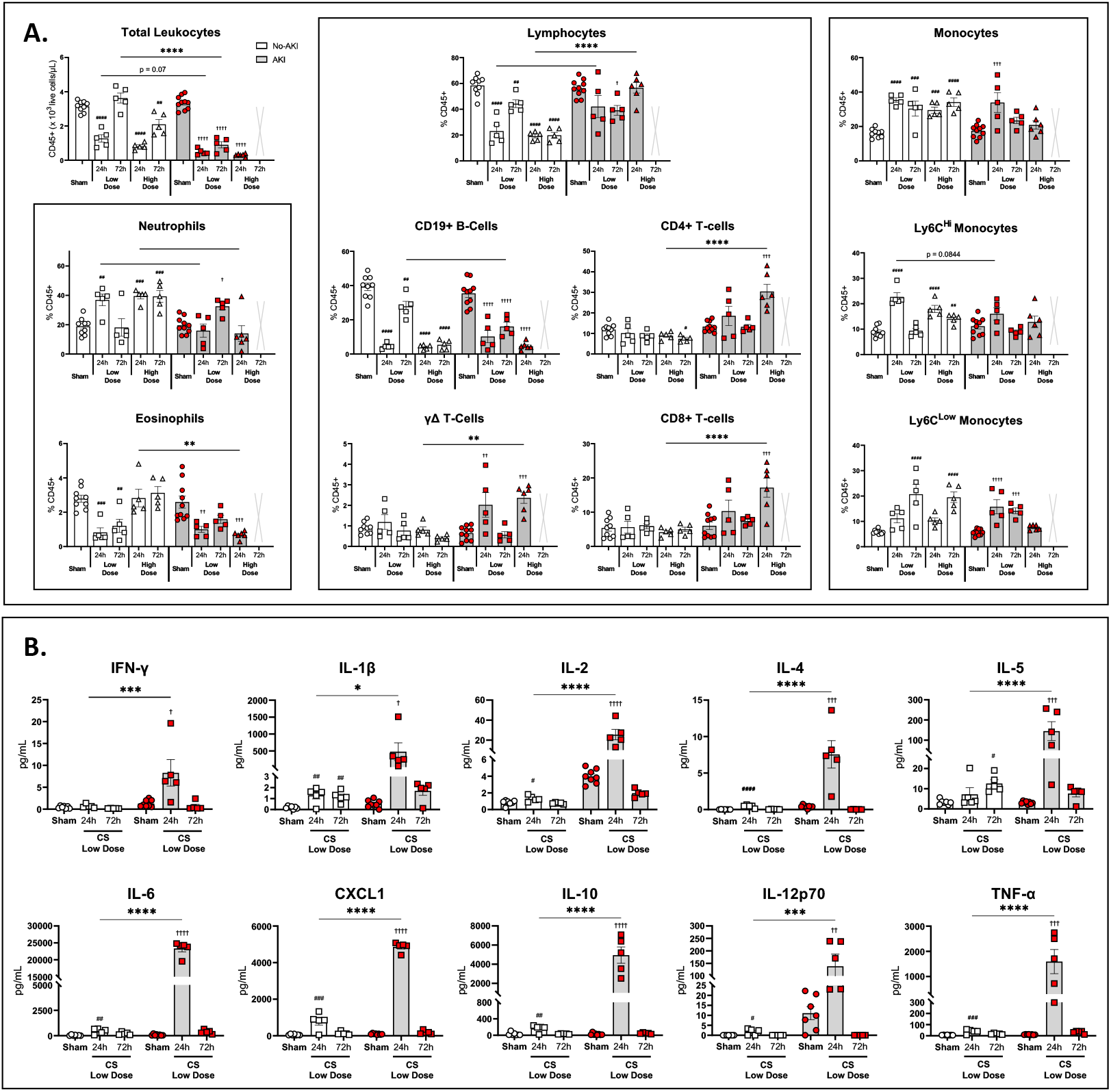
Mice with AKI experience leukopenia, multiple peripheral leukocyte abnormalities, and altered pro-inflammatory cytokine production during sepsis. A) Flow cytometry of peripheral blood leukocytes was performed on mice in the No-AKI and AKI groups at 24- and 72-hours after inoculation (24/72-hour samples are pooled in sham animals). Leukocytes dropped acutely after sepsis then rebound in total numbers in mice without AKI. Conversely, mice with AKI experienced significant leukopenia with an inability to reconstitute total numbers among surviving mice at 72-hours. Among CD45+ leukocytes, individual populations of immune cells were profiled including: granulocytes (A, left panel), lymphocytes (A, middle panel), and monocytes (A, right panel). Compared to No-AKI animals, animals with AKI experienced significant neutropenia and eosinophilopenia, an increase in T-cells, and unchanged to slightly diminished monocyte differentiation in a dose dependent manner. B) Plasma cytokine analysis using a Mesoscale V-plex assay was performed in sham, low-dose 24h, and low-dose 72h samples from No-AKI controls and AKI animals. All profiled cytokines (IFN-γ, IL-1β, IL-2, IL-4, IL-5,, IL-6, CXCL1, IL-10, IL-12p70, and TNF-⍺ were significantly increased in mice with AKI at 24-hours after low-dose sepsis, indicating an exaggerated acute cytokine response, which returned to levels similar to controls by 72-hours after sepsis. (Within No-AKI group: # p<0.05, ##p<0.01, ###p<0.001, ####p<0.0001; Within AKI group: †p<0.05, ††p<0.01, †††p<0.001, ††††p<0.0001; No-AKI vs AKI group: * p<0.05, **p<0.01, ***p<0.001, ****p<0.0001)

Interestingly, neutrophils increased in no-AKI mice at 24-hours after CS inoculation, then fell by 72-hours. Opposite dynamics were appreciated AKI mice (**Figure 5A, left panel**). Eosinophilopenia was appreciated in a dose-dependent fashion similar to dynamics in B-cells. In mice without AKI, evidence of an acute increase in “M1-like” Ly6C-Hi expressing monocytes predominated early, followed by increasing “M2-like” Ly6C-Low monocytes later in sepsis (**Figure 5A, right panel**). Mice with AKI had impaired peripheral monocyte differentiation and, as with other leukocytes, trended towards cytopenias more than No-AKI counterparts.

In order to examine if mice with AKI inoculated with lower dose CS, who had more subtle clinical and biochemical worsening of sepsis severity vs No-AKI mice, had evidence of immune dysregulation-a multiplex cytokine array was performed on peripheral plasma samples (**Figure 5B**). No-AKI animals demonstrated modest, but significant increases in cytokines by 24-hours after CS injection vs sham controls (IL-1β: 1.3 vs 0.2 pg/mL p=0.004, IL-2: 1.3 vs 0.8 pg/mL p=0.02, IL-4: 0.45 vs 0.008 pg/mL, IL-6: 456 vs 40 pg/mL, p=0.002, CXCL1: 795 vs 58 pg/mL, p=0.0005, IL-10: 168 vs 30 pg/mL, p=0.001, IL-12p70: 1.69 vs 0 pg/mL, p=0.01, TNF-α: 39.4 vs 6.8 pg/mL, p=0.0004, respectively). In comparison, AKI animals with low dose sepsis were observed to have massive increases in all cytokines profiled 24-hours after CS inoculation vs AKI sham animals, and vs corresponding No-AKI animals, which largely returned to near normal values in mice collected at 72 hours.

### Adjustment of cecal slurry dose for body weight does not correct differences in sepsis severity between AKI and control mice

Mice with preexisting AKI have lower body weights than controls (**Figures 1B & 3A**). Therefore, using fixed volume doses of CS the AKI mice receive a higher dose-to-body weight than control mice, potentially skewing sepsis severity worse in the AKI mice. To examine if body-weight adjustment in CS dosing would reverse the differences in sepsis severity between AKI and No-AKI controls, separate groups of No-AKI (N=10) and AKI (N=10) mice were generated and received 10ml CS per kg body weight (1mg stool / 1g body weight). Mean weights were 27.8g in No-AKI mice and 22.8g in AKI mice, resulting in CS dosing that was slightly below the 0.3mL high CS dose strategy in No-AKI mice, and CS doses slightly above the 0.2mL low-dose strategy in AKI mice (**Supplemental Figure 3A**). Body weight correction did not result in narrowing of comparative sepsis severity in No-AKI and AKI mice. And despite only slightly higher delivered CS dose to AKI mice vs low-flat CS dose strategy, these animals experienced significantly more severe sepsis and rapid animal attrition, while No-AKI mice again experienced minimal distress (**Supplemental Figure 3B, 3C**).

### Mice with preexisting CKD experience less dramatic increases in sepsis severity versus controls without CKD

Generation of No-CKD (N=30) and CKD (N=30) mice proceeded as previously outlined (**Figure 1A**), and the CKD donor stool collection and experimental details was performed as in AKI studies (**Figure 2A-C**). Baseline weight and tGFR measurements are displayed in **Figure 6A** and **6B**. Similar to studies in **Figure 1**, the No-CKD control mice differed from No-AKI control mice only in duration of vehicle-injections, age, and body weight. As with No-AKI controls, the No-CKD control group experienced no attrition with either low or higher CS doses, with similar dynamic trends of temperatures and sepsis scores throughout infection evolution (**Figure 6C-G**). In contrast to No-AKI controls, No-CKD controls lost considerably more weight with either low or higher CS doses at 24h (−8.8% and −11.1% weight loss, respectively) and failed to regain weight by 72 hours (−11.1% and −13.4% weight loss, respectively). This may be a consequence of the older age of mice, or higher baseline body weight vs the No-AKI controls before CS studies. Mice with CKD tolerated CS injections better than mice with AKI at either flat-dosing CS strategy, although two animals were euthanized (one at 36 hours and another at 48-hours, according to criteria) in the 72-hour high-dose studies. Surviving mice with CKD had similar weight changes after CS inoculation compared with No-CKD mice, with a statistically insignificant trend towards less overall weight loss.

**Figure 6.**
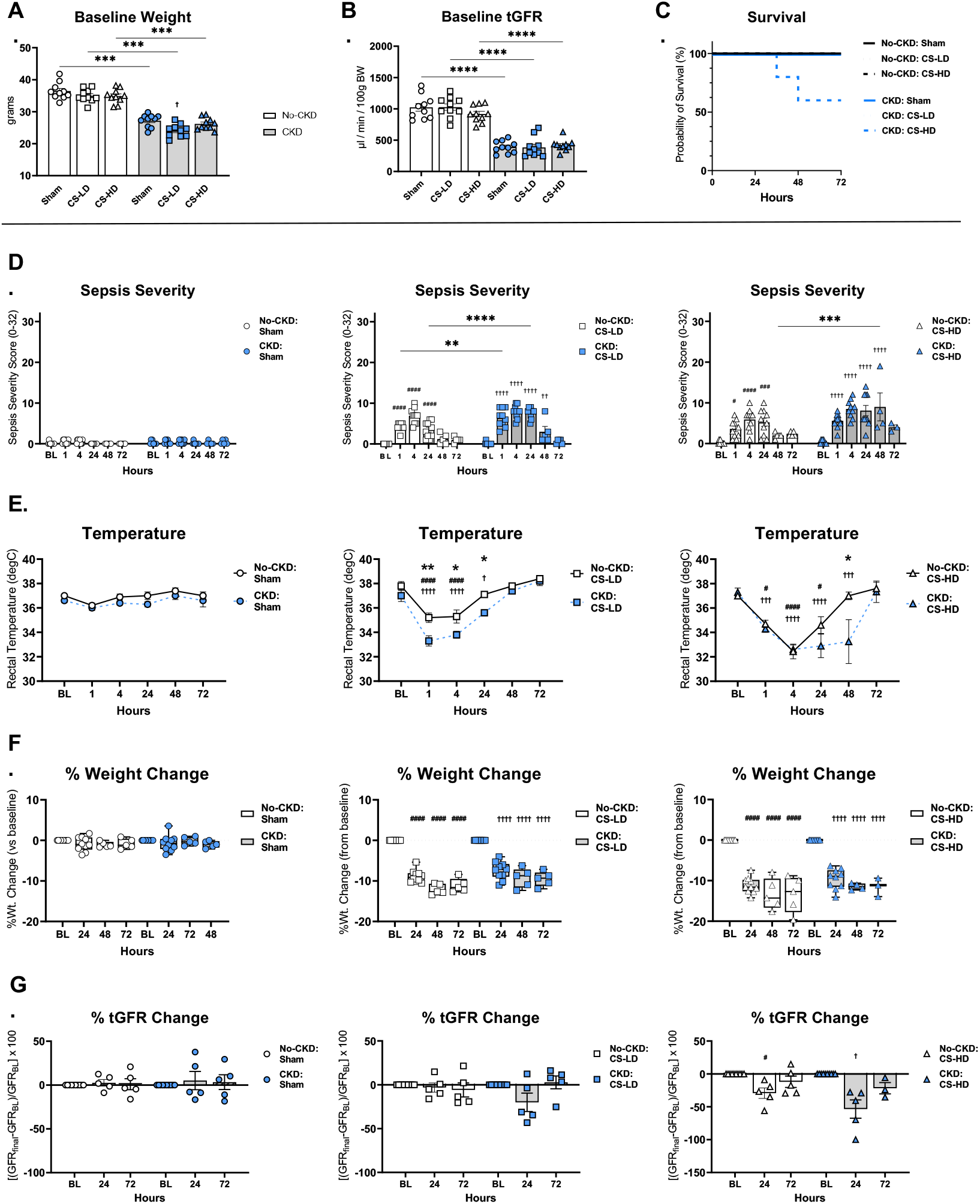
CKD CS Sepsis Studies. A) Baseline weights and B) tGFRs are again shown in No-CKD control (N=30) and CKD mice (N=30). C) Two CKD mice receiving CS-HD required euthanasia: occurring at 36-, and 48-hours after inoculation. As in AKI studies, control animals had 100% survival in all groups. D) Sepsis severity, E) rectal temperatures, F) change in weight vs baseline, and G) change in GFR vs baseline are shown. With both low-and high-CS doses CKD mice experienced significantly worse sepsis severity and hypothermia, though trends in weight and GFR loss were similar to controls. (Within No-CKD group: # p<0.05, ##p<0.01, ###p<0.001, ####p<0.0001; Within CKD group: †p<0.05, ††p<0.01, †††p<0.001, ††††p<0.0001; No-CKD vs CKD group: * p<0.05, **p<0.01, ***p<0.001, ****p<0.0001)

tGFR measurements in sham animals from either group showed no significant changes across the study (**Figure 6G**, **Supplemental Figure 2B**). With low-or higher CS doses, No-CKD and CKD mice showed similar patterns of renal functional decline; with only higher CS doses inducing statistically significant decline in tGFR in both 24-hour groups, though the magnitude of decline from baseline was not significantly different when comparing the two groups (**Figure 6G**). As in AKI studies, surviving mice at 72 hours in both groups experienced return of kidney function near baseline. Bacteremia was again assessed, and two CKD mice (the animals with most severe sepsis requiring euthanasia at 36- and 48-hours) had evidence of detectable bacteremia (**Supplemental Figure 1A**).

Plasma measurements of BUN, AST, and NGAL showed similar overall dynamics in the CKD CS studies as appreciated in AKI studies, with some notable exceptions (**Figure 7A, 7B**). Baseline levels of BUN were higher in CKD mice than No-CKD mice (49 vs 24 mg/dL, p=0.005, respectively). As an overall sepsis biomarker, AST again performed reasonably well, peaking at 24-hours after high-dose sepsis in No-CKD mice (71 vs 26 U/L, p<0.0001), and in CKD mice (90.8 vs 24 U/L, p<0.0001); with CKD mice having significantly higher levels than No CKD mice. Plasma NGAL again performed well as a sensitive and specific sepsis biomarker, mirroring the clinical data in the CKD group the strongest. Acute tubular injury was noted in a dose-dependent manner in No-CKD control mice (**Figure 7D**), again appearing to peak in a delayed fashion 72-hours after injury. As with AKI specimens, kidneys from CKD mice had severe baseline tubular injury, and determining any new changes in new tubular injury vs sham-injected animals was difficult by histological methods. Objectively more proteinaceous cast formation was appreciated in kidneys from septic CKD mice, though this finding was not statistically significant (**Figure 7C**).

**Figure 7.**
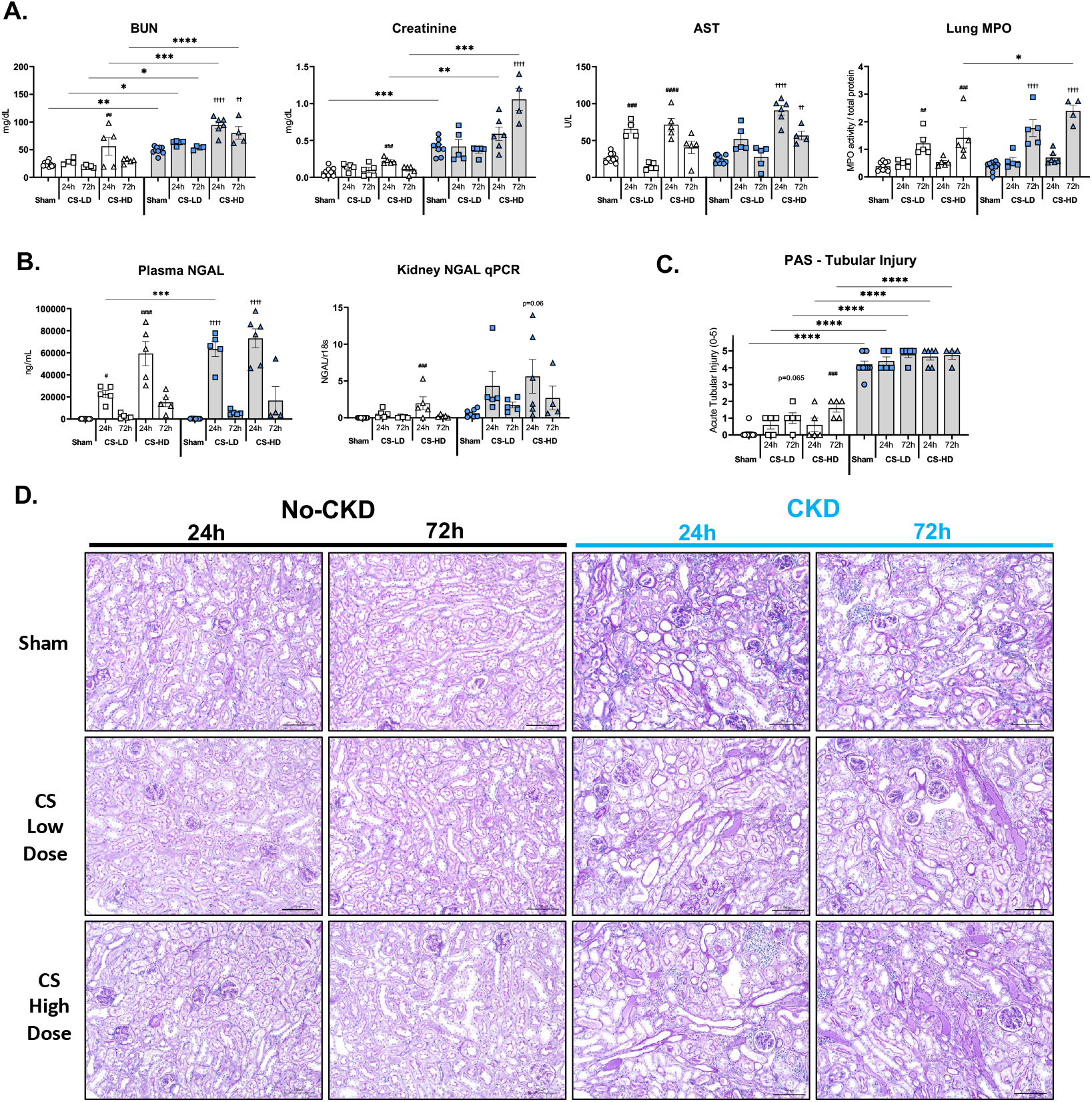
Mice with CKD have worse organ injury than No-CKD controls after sepsis. A) Plasma BUN, creatinine, AST, and lung MPO levels were determined. At baseline, mice with CKD had significantly higher plasma BUN and creatinine than control mice. AST values mirrored plasma and kidney NGAL levels (B), peaking at 24h in the model with plasma NGAL showing better sensitivity. B,C) PAS-stained cortical tissues from no-CKD controls showed significantly increased tubular injury by 72 hours in both low and high-dose CS studies. However, due to the extreme baseline tubular injury in CKD mice, only a minor, non-significant trend towards increased tubular injury was appreciated in higher CS dose studies, driven primarily by the appearance of more proteinaceous tubular casts vs. controls. (Within No-CKD group: # p<0.05, ##p<0.01, ###p<0.001, ####p<0.0001; Within CKD group: †p<0.05, ††p<0.01, †††p<0.001, ††††p<0.0001; No-CKD vs CKD group: * p<0.05, **p<0.01, ***p<0.001, ****p<0.0001)

### CKD mice have isolated, modest peripheral immune defects and cytokine elevation after CS inoculation vs No-CKD controls

Unlike AKI studies, mice with CKD had less severe differences in flow cytometry-sorted populations of leukocytes vs No CKD controls. Patterns of absolute leukopenia after CS injection were similar in both groups as compared to AKI studies, with CKD mice showing non-significantly lower total leukocyte blood numbers at 72 hours in both low and high dose CS studies (**Figure 8A**). CKD mice demonstrated significantly lower neutrophils, eosinophils, and significantly higher T-cell populations vs No-CKD mice, but these differences were overall modest. No differences were determined in peripheral blood mononuclear cell (PBMC) differentiation in infected CKD vs No-CKD control mice. Plasma cytokines profiled from animal groups receiving low dose CS were again measured (**Figure 8B**), and CKD mice showed significant increases in IL-2, IL-6, CXCL1, IL-10, IL-12p70, and TNF-α vs No CKD control mice at 24-hours, though absolute increases at this time point, and with this CS dose were comparatively lower than those observed in infected AKI mice. Interestingly, IL-1β and IL-5 levels were significantly higher in infected controls vs CKD mice at 72 hours. The relative increase in IL-5 levels in control mice was accompanied by higher eosinophil levels profiled by flow cytometry at this time point (**Figure 8A, left panel**).

**Figure 8.**
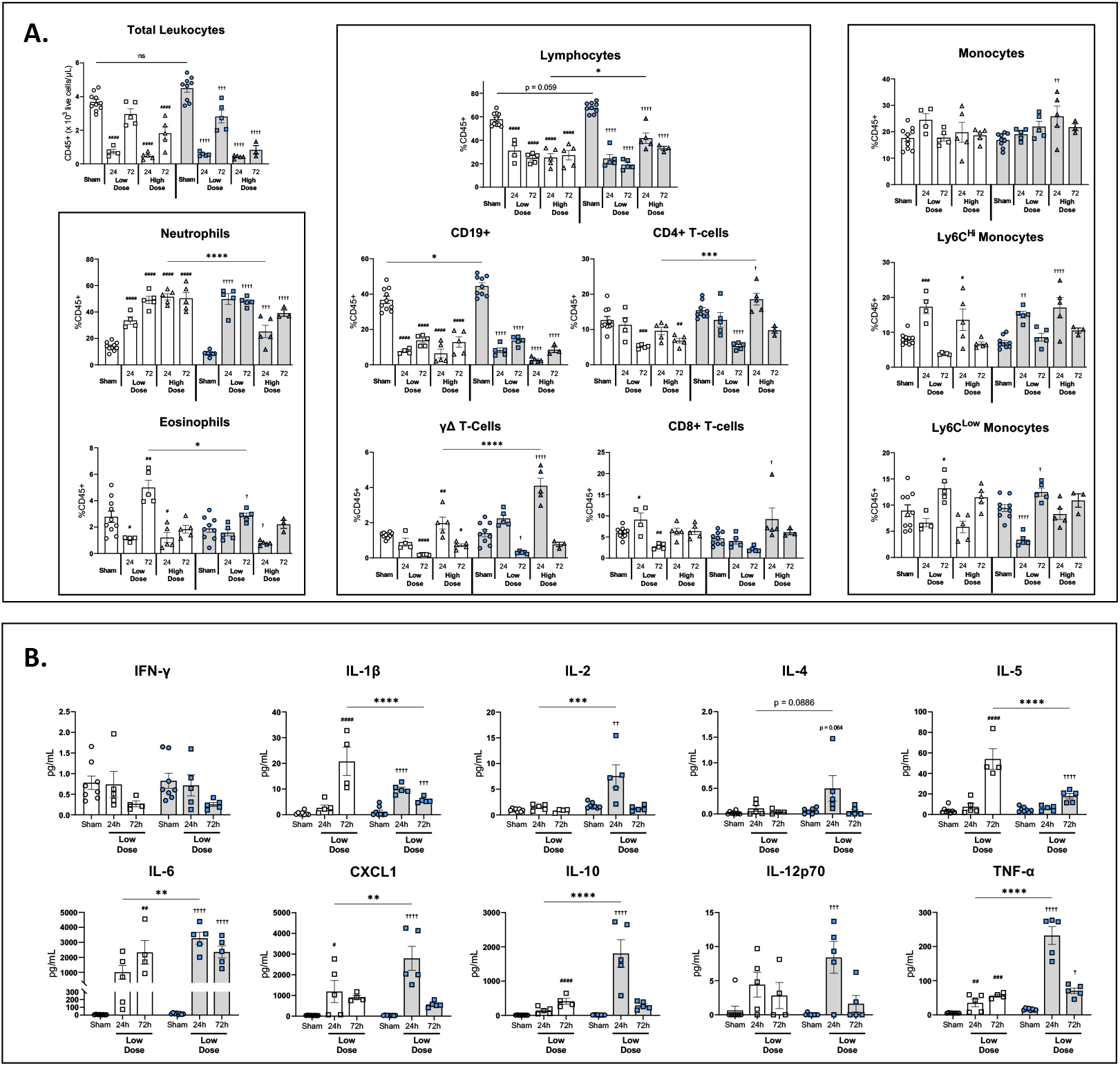
Mice with CKD have a similar peripheral blood immune response to sepsis vs control mice. (A) Flow cytometry of peripheral blood leukocytes was performed on mice in the No-CKD and CKD groups. 24- and 72-hour sham samples are shown pooled as before. Populations are expressed as percent CD45+ cells. CKD mice had similar degrees of leukopenia after sepsis compared to control animals (A, upper left panel). Aside from a significant increase in CD4 and γΔ T-cells, and significant decrease in neutrophils in high-dose sepsis 24h studies, populations of profiled immune cells from CKD mice largely mirrored no-CKD controls. (B) Plasma cytokine analysis using Mesoscale Vplex assay. Mice with CKD were observed to have significantly higher IL-2, IL-6, CXCL1, IL-10, and TNF-⍺ levels vs controls at 24-hours after low-dose sepsis. Interestingly, mice with CKD had blunted IL-1β and IL-5 levels by 72-hours after low-dose sepsis vs controls. (Within No-CKD group: # p<0.05, ##p<0.01, ###p<0.001, ####p<0.0001; Within CKD group: †p<0.05, ††p<0.01, †††p<0.001, ††††p<0.0001; No-CKD vs CKD group: * p<0.05, **p<0.01, ***p<0.001, ****p<0.0001)

### Body weight CS dose adjustment partially corrects sepsis severity between CKD and control mice

As in AKI studies, CKD mice and controls have divergent weights versus controls (**Figures 1C, 6A**), potentially leading to inappropriate skewing of sepsis severity in CKD mice if using a flat CS dosing strategy. Body weight adjustment experiments were again performed at 10 ml/kg CS-to-body weight. Mean body weights at time of inoculation were 34.4 and 27.7g, resulting in an administration of cecal slurry doses that were higher in No-CKD mice than the 0.3 mL flat-high dose experiments, and dosing slightly below the 0.3 mL flat-high dose experiments in CKD mice (**Supplemental Figure 3D**). Body weight adjustment partially corrected sepsis severity between No-CKD and CKD mice. CKD mice continued to experience moderate later attrition of numbers, though importantly two animals in the No-CKD control group required euthanasia (**Supplemental Figures 3E, 3**). These were the only control animals with preexisting normal kidney function in all the studies that did not survive CS injections.

Of note, these two euthanized animals in the control group were also the animals receiving the highest CS doses given their baseline weights, which were comparatively heavier than littermates (40.1 g and 37.9 g vs mean of 33.3 g in the other littermates). The heaviest dosed mouse in the CKD group (31.0 g) was also one that experienced the highest sepsis severity. Among other CKD mice there was no relationship between dose or sepsis severity, though there was little variability in baseline weights, and therefore administered CS doses (standard deviation of 1.39 g body weight in other N=9 CKD animals). If the two heaviest control mice were excluded from analysis, sepsis severity would appear similar to the flat-high CS dosed No-CKD control group. These data indicate that a weight adjustment dosing strategy might cause *more* variability in sepsis response, especially if: 1) there is a large standard deviation of weights between animals in the same experimental group, and 2) adjustments are made at relatively higher CS doses, since effects of CS appear to work in a semi-linear dose-response manner, until a certain threshold is reached then sepsis severity appears to progress in a non-linear relationship with dose.

## Discussion

We have described a new parallel method of creating AKI and CKD with repeated, low-dose injections of aristolochic acid; which was then utilized to study the dynamics of acute intrabdominal infection using donor-recipient matched CS inoculation. We believe these studies are important for several key reasons. 1) The AA model, while having some variability, nevertheless appears to be well suited to reliably reproduce clinically stable states of AKI and CKD that can be used for comparative studies. 2) The CS live-bacterial infection approach allows for a study of acute infection in a “dose-dependent” manner that may not be possible with more blunt methods like surgical CLP given the fragility of the underlying host. 3) We have shown that AKI predisposes to worse infection outcomes than CKD; and this observation occurred despite higher baseline measured GFR in mice with AKI, and at younger animal ages. 4) Finally, we have demonstrated that “AKI-on-AKI” and “AKI-on-CKD” can be effectively modeled and monitored non-invasively using serial assessments of measured GFR. Therefore, our findings may have translational implications beyond the immediate scope of this study.

Our data add to the literature regarding the dysregulated nature of immunity in hosts with underlying AKI and CKD. While there are many observations of the nature of immune defects imparted by AKI and CKD in humans, there is little preclinical data that can model this unique host-pathogen interaction. We are aware of one group of investigators who successfully performed CLP in the setting of both AKI induced by acute folic acid (FA) administration,^26^ and CKD induced by remnant nephrectomy.^27^ In these studies both AKI and CKD predisposed to severe early morbidity and death versus control CLP mice, with few animals surviving past the first 24-hours post CLP.

We believe that using CS to induce sepsis in these uniquely immunocompromised hosts allows for a more nuanced study of both the absolute severity of infection, but also the different phases of infection and recovery. Animals with AKI experienced a hyperacute inflammatory response, yet had early attrition with high degrees of bacteremia, and severe cytopenias. We hypothesize that these animals developed rapid septic shock, as a similar massive acute peripheral cytokine response has been observed in humans with the highest degrees of sepsis severity.^28, 29^ AKI animals interestingly had evidence of a simultaneously upregulated acute (IFN-γ, IL- 1β, IL-6, TNF-α) and counter-acute inflammatory response (CARS, IL-10 meditated) cytokines, which has also been appreciated in humans with severe sepsis.^30^ A distinguishing feature in AKI mice with low and high doser of CS was the severity of peripheral neutropenia and lack of peripheral monocytic differentiation, even in clinically recovered animals. It is possible that early functional exhaustion in these cells resulted in poor pathogen clearance leading to rapid septicemia and death.

In CKD mice there was more delayed morbidity with CS injection versus controls, and peripheral leukocyte dynamics during sepsis evolution followed a similar overall pattern as appreciated in control mice. In both AKI and CKD mice, significant B-cell depletion and T-cell activation was appreciated, suggesting similar adaptive immune defects might be present. The most surprising observation from our studies was that our CKD mice, having much lower baseline tGFR, older age, and (historically appreciated) much more severe baseline anemia, had less acute (t=0 to t=24h) sepsis severity and animal attrition at any dose of CS vs AKI mice.

One simplistic interpretation of our studies is that mice with AKI and CKD might simply have a lowered threshold for severe sepsis from CS peritonitis. However, among the surviving AKI or CKD mice that received lower CS doses there was evidence of poorer overall infection recovery (sepsis score, temperatures, weight gain, and peripheral reconstitution of leukocytes) versus respective control mice receiving the highest CS doses (who had roughly matched t=1h, 4h, 24h sepsis scores and temperatures). These data indicate that mice with preexisting kidney disease may also have a delayed defect in infection resolution that is not explained by degree of acute (clinically) determined sepsis severity.

The donor-recipient matching of CS was used to mimic an endogenous source of host infection, similar to what would occur in the setting of a perforated intestinal viscus. We cannot determine from our data if intentionally mismatching donor CS-recipient groups would change the clinical response to infection. Such studies might determine if divergences in sepsis severity are more likely due to preexisting host immunodeficiency *versus* unique differences in the host microbiome. With regard to the CS vehicle solution, we followed published methods^21, 22^ suspending the CS homogenates in 10% glycerol in PBS. This method has been shown to maintain CS virulence with no loss in numbers of bacteria grown after prolonged (> 6 months) periods of storage. Pertinent to our studies, there is an appreciation that ATI can accompany high loads (30- 50%) of glycerol administration, ^31^ from the creation of spontaneous rhabdomyolysis. We found no appreciable increase in ATI in control, AKI, or CKD mice receiving sham PBS-glycerol injections assessed by histology, biochemically with BUN, Cr, as well as directly measured GFR. We administered higher doses of PBS-glycerol (0.3mL) to all sham animals to mirror the highest possible dose of PBS-glycerol the infected animals were given in anticipation of this potential confounder. Additionally, PBS/glycerol-injected AKI mice actually demonstrated improvement in renal function, which we ascribe to expected AKI recovery soon after the cessation of AA-injections.

Our study has some limitations. Some physiological data of sepsis severity is lacking; and tGFR measurements were performed only at baseline and at terminal collection. Given the degree of frailty in the animals after CS inoculation, we chose to manipulate them as infrequently as possible so as not to influence their clinical outcome. Also, we did not to perform baseline, or serial blood draws on individual animals given that this intervention might also influence sepsis severity. This concern was especially relevant among animals with preexisting kidney injury, who have evidence of significant anemia at baseline. Separate sham-operated animals were used as alternative controls in this regard. We did not administer prophylactic fluids (except in a reactive manner with severe sepsis as outlined in methods), nor antibiotics so cannot determine if the mice with high morbidity and mortality would be rescued with these interventions.

We chose 24- and 72-hours as short and medium-term clinical endpoints for the model based on published data and our own observations of sepsis evolution after this method of CS injection. We acknowledge that our data on plasma cytokines, flow cytometry, and measured GFR are not a complete picture-that understanding much earlier events, such as occurring at 1-4-hours after inoculation (when earlier evidence of clinical divergence in animal groups was first observed) is important to give more context to our studies. Additionally, clinical responses to CS inoculation should be assessed in surviving mice beyond 72-hours, particularly given our observations that immune dysregulation in surviving animals with kidney disease is persistent at this later time point. Our AKI and CKD groups were also not age matched at baseline, with a modest difference of 9 weeks between the groups immediately before sepsis studies commenced. Others have reported an age-related intolerance to CS injection^21^, therefore it is possible that aging our AKI mice to match the CKD animals would have resulted in even worse infectious outcomes.

The AA-model itself might influence the sepsis phenotype, independent of degree of kidney injury and measured GFR. AA has been associated with the development of anemia^32, 33^, findings confirmed in our initial studies. However, AA is not known to induce other immune cell defects; and we did not appreciate that AKI or CKD mice had significantly different peripheral immune cell populations versus controls at baseline. We cannot determine if a unique bone-marrow myeloid differentiation defect is more present in the AKI mice. However, the fact that CKD mice, with lower baseline hemoglobin concentrations and more cumulative AA exposure, had relatively intact peripheral reconstitution of leukocytes after CS infection makes any AA-specific bone marrow defect unlikely. The degree of baseline anemia tracked overall with renal mass and GFR loss in the model, as one might expect in humans with progressive CKD.

Finally, we acknowledge that our findings might be different with alterations in animal strain, species, sex, and different models of kidney injury. In our studies, the AKI model was designed to mimic moderate ATI with depression in GFR for weeks, rather than an abrupt insult and recovery, such as occurs in ischemia/reperfusion (I/R). This method was chosen to maximize AKI survival in animals and minimize variability in degree of baseline kidney injury prior to CS inoculation. Our CKD model effectively mimicked many renal and extrarenal manifestations of human CKD with high fidelity, as these animals had prolonged exposure to GFR decline rather than a sudden reduction in nephron mass and GFR that occurs with surgical models like remnant nephrectomy. We determined that using a nephrotoxic model like AA-nephropathy was ideal for our studies, as it potentially minimizes any preexisting priming of peritoneal immune cells that could occur with kidney injury models requiring prior abdominal surgery.

In conclusion, we have demonstrated that clinically relevant AKI and CKD clinical states can be modeled in a rodent with slow aristolochic acid injections, which is a suitable background model to then deploy comparative studies of sepsis using the CS method of peritoneal infection. We believe this novel model can serve as a backbone to investigate host immune defects in AKI and CKD with more rigor; and may aid in the testing of interventions designed to improve clinical outcomes in sepsis. Furthermore, in our studies we observed that the phenotype of kidney disease present at baseline (AKI versus CKD) appears to influence sepsis severity more than degree of baseline renal failure as measured by GFR. Immunodysregulation in the context of differing clinical state of kidney disease should resultingly be a high priority area for further research.

## Methods

### Rodent model of Acute and Chronic Aristolochic Acid Nephropathy (AAN)

All animal studies were approved by the Institutional Animal Care and Use Committee at the University of Colorado Anschutz Medical Campus and the Rocky Mountain Regional VA Medical Center. C57BL/6J male mice were procured and underwent experimentation starting at 8-10 weeks of age. Vehicle control (phosphate buffered saline, PBS) or AA (AA-1 sodium salt, Sigma) was administered at a dose of 2 mg/kg intraperitoneal (i.p.) thrice weekly x 3 week for the AKI model, and thrice weekly x 3 weeks plus weekly x 3 weeks, then allowed to recover for an additional six weeks for the CKD model (**Figure 1A**). Animals used for the initial model description underwent serial assessments of weight and transcutaneous glomerular filtration rate (tGFR, see below) at baseline, and weekly until euthanasia in the AKI model, and baseline followed by assessment at 3, 6, and 12 weeks in the CKD model. These animals were euthanized on day 21 (AKI model) or day 84 (CKD model). Separate animals used for the cecal slurry (CS) sepsis studies underwent weight and tGFR assessment only immediately prior to inoculation, after kidney disease had been established.

### Transcutaneous GFR

tGFRs were assessed for animals in **Figure 1** as previously mentioned. For CLP pilot studies tGFRs were assessed at baseline only (day before cecal ligation and puncture). For CS sepsis studies, all measurements were taken on days 21 or 84: (1) as a baseline measurement and (2) 24- or 72-hours after sham or cecal slurry inoculation. Mice were shaved 1-2 days prior to initial baseline measurement to reduce skin aggravation on the day of device readings. Animals were anesthetized with 1.5-2.5% isoflurane, placed on a heating platform (Kent Scientific), where the shaved area on the right dorsal thorax was cleaned with 70% ethanol and allowed to dry. A charged battery was attached to the tGFR-mini device (Medibeacon, Mannheim Germany), which was then attached to the animals with a proprietary transdermal sticky pad. Surgical tape was used to snugly secure the device in place. FITC-Sinistrin (10mg/100g body weight, Medibeacon) was then administered via a retro-orbital injection and mice monitored for a minimum of 70 minutes. Measurements were read with proprietary software and GFR derived from FITC-Sinistrin half-life as per the manufacturers recommendations^34^ and as detailed in our previous publications.^35–37^

### Cecal Slurry Preparation

Cecal slurry creation was performed as described originally ^20^, with important recent modifications ^21, 22^. Sham-sepsis animals (see procedure below) were identified as suitable cecal slurry donors and, at the time of terminal collection, underwent isolation of cecum, sterile dissection of contents, suspension of contents into 10% glycerol in sterile PBS at 100mg stool/mL (**Figure 2B**). Contents were press-filtered serially through sterile 860, 320, 180µM filters (Bellco Glass, Inc., Vineland, NJ) and a 100µM sterile cell strainer (ThermoFisher Scientific, Waltham, MA). A small aliquot (50-75µL) of the final solution from each animal was separated for later 16s DNA analysis. The final solution from each animal was combined with similar grouped littermates to create a homogenous host-matched cecal slurry. Slurry was then aliquoted into separate cryovials under conditions of continuous stirring using a sterile magnetic bar on stir plate, frozen on dry ice immediately and stored at −80degC. Cecal slurry stocks were used within 6 weeks of generation, though published data indicate bacterial loads are completely stable up to 6 months and beyond using this method^21^. On day of experimentation, slurry vials were rapidly thawed in a 37degC shaking water bath, transported to the vivarium and kept warm on a 37degC heat block for a short period until injection. Shortly before injection, vials were agitated by hand, sample drawn up with a sterile 0.5mL tuberculin syringe and injected into the right lower ventral abdomen into the peritoneal space at the indicated volume.

### Cecal Slurry Peritonitis

Newly generated AKI-and CKD-mice underwent cecal slurry peritonitis experimentation starting at the two previously established AAN time points (**Figure 2A**). Each AKI-and CKD-cohort was accompanied by separate vehicle injected animals as before, designated as “No-AKI” and “No-CKD” controls. No-AKI, AKI, No-CKD, and CKD mice received either a single intraperitoneal injection of sham-vehicle solution (10% glycerol in PBS at 0.3mL), low dose (0.2mL, 20mg), or high dose (0.3mL, 30mg) host-matched cecal slurry. Additional groups were generated for the supplemental body weight dose-adjustment experiments (10ml CS/kg body weight, or 1mg/g, **Supplemental Figure 3**). Within each dosing strategy, separate paired groups were generated for planned analysis at either 24- or 72-hours after inoculation. Mice were euthanized after final tGFR performed via terminal isoflurane inhalation using a drop method, animals dissected with bilateral thoracotomy and laparotomy, EDTA-whole blood obtained by cardiac puncture and separated for analysis of frozen plasma or flow cytometry; kidneys and lungs dissected and placed in 4% paraformaldehyde for histology and separate tissues frozen on dry ice.

### Sepsis scores, temperatures, and weight changes

#### CLP Pilot Studies

a simplified sepsis severity score was created for assessment at t=24h in surviving animals (0=no distress, 1=mildly limited spontaneous movement, responds to auditory stimulus, mild/moderate piloerection of coat, 2=moderately limited spontaneous movement, responds to auditory stimulus, mild/moderate piloerection of coat, 3=moderately limited spontaneous movement, poorly responds to auditory stimulus, moderate/severe piloerection of coat, 4=severely limited spontaneous movement, poorly responds to auditory stimulus, moderate/severe piloerection of coat, 5=moribund with no spontaneous movement, no response to auditory stimulus, moderate/severe piloerection of coat, may have rigors. Weight changes were also assessed at t=24h collection vs baseline (before surgery).

#### CS Peritonitis Studies

For CS studies the sepsis score was extensively expanded using the framework published by Shrum B et al.^38^) with slight modifications (**Supplemental Figure 4**). Additionally, the score was obtained serially alongside rectal temperature probe measurements (PhysioSuite, Kent Scientific, Torrington, CT) at: baseline and hours 1, 4, 24 in all animals; and at 48 and 72 hours in the 72-hour designated groups. Weight changes were assessed at baseline, and daily until planned collection.

### Cytokine analysis, organ injury, bacteremia

Plasma was isolated from EDTA-anticoagulated whole blood, centrifuged at 2,000 xg for 10 minutes at 4degC, and frozen at −80degC for future measurements. Blood urea nitrogen (BUN), Creatinine, and Aspartate transaminase (AST) were assessed using commercially available kits (BioAssay Systems, Hayward CA). Lipocalin-2 (NGAL) was assessed with a Quantikine ELISA Kit (Biotechne R&D Systems, Minneapolis MN). Additionally, a multiplex inflammatory cytokine panel (IFN-γ, IL-1β, IL-2, IL-4, IL-5, IL-6, IL-10, IL-12p70, KC/GRO (CXCL1), and TNF-α, MesoScale Discovery, Mouse V-Plex Proinflammatory Panel 1) was performed following manufacturer instructions and analyzed using the MESO Quickplex SQ120 instrument and Discovery

Workbench software. Lung myeloperoxidase (MPO) activity was measured with frozen harvested tissue. Lung samples (30-50mg) were suspended with 1mL of HTAB buffer (2g hexadecyltrimethylammonium bromide in 400mL MPO Buffer-3.4g of potassium dihydrogen phosphate and 4.35g of dipotassium hydrogen phosphate in 450mL deionized water, pH 6.0) and homogenized using a sterile 5mm stainless steel bead in a Tissue Lyser 2 (Qiagen, Hilden, Germany) for 90 seconds at 1/60Hz. The homogenate was transferred to a 2mL tube and sonicated for 10 seconds, then centrifuged at 14,000 xg for 30 minutes at 4degC. MPO activity was measured by combining supernatant with 3% hydrogen peroxide in o-dianisidine MPO solution (16.7mg o-dianisidine in 90mL DI water with 10mL MPO buffer, heated to 37degC). A plate reader was used to measure absorbance at 450nm. Total protein quantity was measured using a Bradford Assay Kit (ThermoFisher Scientific). Bacteremia was assessed by plating 10µL of EDTA-whole blood onto trypsic soy agar plates, serial diluted 1:2, then by a factor of 10 to 1:1000, incubated for 24 hours at 37degC, and detection was determined. Of note in several of the high grade bacteremia samples we were unable to quantify number of colonies as they were not diluted enough, therefore results were reported as positive/negative rather than determination of CFU counts due to incomplete data.

### Histology for Acute Tubular Injury and Fibrosis

Harvested mouse organs (kidney, lung) were preserved in 4% paraformaldehyde (Electron Microscopy Sciences, Hatfield, PA) for 24 hours before being processed and embedded in paraffin-blocks. Blocks were cut to 5µM sections and stained for collagen using Picrosirius Red Staining (Model development studies only-**Figure 1**); and fibrosis was quantified using an analysis of polarized PSR-stained cortical tissues as we have previously outlined^39^. Separate sections were stained using a commercially available periodic acid Schiff (PAS) staining kit (Newcomer Supply, Middleton, WI). Acute tubular injury was scored on PAS stained sections (Model development, AKI, and CKD studies) using a modification of published methods^40, 41^ as follows: Injury including evidence of brush border loss, tubular dilation or atrophy, cell necrosis, presence of proteinaceous casts, and degree of tubulitis and interstitial nephritis was graded from 0-5 with none (0), < 10% (1), 11-25% (2), 26-45% (3), 46-75% (4), and > 76% (5) of affected corticomedullary kidney regions. If corticomedullary boundary could not be determined (as in CKD samples), random mid-cortical regions were analyzed. Approximately 3 separate sections of transverse cut kidney tissues were examined with 4-6 20x light microscopic fields examined per section and scored in blinded fashion, with sample identification masked. Samples were randomly scored a second time and an average of the two scores were obtained.

### Peripheral Blood Flow Cytometry

EDTA-treated whole blood obtained by cardiac puncture (150µL) was treated with RBC lysis buffer (eBioscience, ThermoFisher Scientific) for 3 minutes, washed with FA3 buffer (PBS, 10mM HEPES, 2mM EDTA, 1% FBS), centrifuged at 300 xg x 5 minutes (underwent an additional RBC lysis for 1 minute and wash/centrifuge again if needed to clear sample), passed through a 70µM filter, and resuspended in FA3 buffer before being plated on 96-well plate for staining. Cells were counted using an automated cell counter (BioRad, Hercules, CA). After block with anti-mouse CD16/CD32 (Invitrogen, ThermoFisher Scientific) and stain with a viability dye (GhostDye Violet 510, Tonobo, Cytek Biosciences, Fremont, CA), cells were incubated at 4degC with the following fluorochrome conjugated primary anti-mouse antibodies for 45 minutes (all Biolegend, San Diego, CA): anti-CD45 PE-CF594 (clone 30-F11), anti-Ly6G PECy7 (clone 1A8), anti-CD11b PE (clone M1/70), anti-Ly6C BV785 (clone HK1.4), anti-TCRβ APC (clone H57-597), anti-CD4 BV605 (clone RM4-5), anti-CD8 APC-Cy7 (clone 53- 6.7), and anti-CD19 PacBlue (clone 6D5). Cells were then centrifuged, washed and resuspended in FA3 buffer, and finally fixed in 2% PFA. Cells were processed through a Cytoflex LX flow cytometer (Beckman Coulter, Indianapolis, IN), and data analyzed using FlowJo single-cell analysis software (FlowJo, Ashland, OR). Gating proceeded as follows: Live è singlets è CD45+ è (CD19+ B-cells) CD19- è (TCRβ+ T-cells è CD4 vs CD8 T-cells (γΔ-Tcells as double negative population)) TCRβ- è (Ly6G+SSC-Hi Neutrophils) Ly6G-SSC-Low è CD11b+ è Ly6C+SSC-low Macrophages, Ly6C-SSC-low Macrophages, Ly6C-intSSC-high Eosinophils.

### Statistics

A student’s T-test was performed for 2 variable comparison, 1-way ANOVA for more than 2 variables, with Tukey’s multiple comparison post-test. For grouped analyses, a 2-way ANOVA was performed with Šídák’s multiple comparisons post-test. For comparisons within groups, statistical significance was only determined between time-and dose-endpoints versus the corresponding sham condition (i.e. sham vs CS-LD 24hr, sham vs CS-LD 72hr), whereas comparisons between different groups included analysis of only parallel conditions (i.e. No-AKI CS-LD 24hr vs AKI CS-LD 24hr). For sepsis studies aggregated clinical data was used for weights, sepsis scores, and temperatures, whereby data from N=10 experimental animals was used at: baseline, 1 hour, 4 hours, and 24 hours after inoculation. Thereafter (48h, 72h) data included only the N=5 remaining animals in the 72-hour designated group. If the comparisons between groups contained missing data because animals did not survive to predetermined endpoint, these data were excluded from analysis. Samples collected prior to predetermined endpoint due to moribund status were included in either 24- or 72-hour analyses on an individual basis. The involved animals included: all AKI animals receiving CS-HD designated for collection at 72 hours (all euthanized at, or shortly before, 24 hours as outlined and were considered as additional 24-hour animals). Additionally, two animals in the CS-HD dose CKD 72-hour studies needed euthanasia before predetermined endpoint; one (moribund by 36 hours) was included with animals in the 24-hour group, and the other (moribund by 48 hours) was included with the 72-hour group in the analysis of biochemical and histologic measurements.

## Acknowledgments

Funding for this study was supported through a Veterans Heath Administration Career Development Award for JRM (VHA BLR&D IK2BX003839). Thanks to Dr. Hiroshi Saito (University of Kentucky) for consultation regarding the cecal slurry model development. Parts of **Figures 1 and 2** created with BioRender.com.

---

**Supplemental Figure 1. Burden of Bacteremia in Septic Mice.** Blood samples were plated on trypsic-soy agar plates to determine bacterial growth. A) Table demonstrating the numbers of animals with positive blood culture plates. (B) Pictures of plates from a group of No-AKI and (C) AKI animals at 24-hour time collection. Undiluted (U) whole blood specimens were plated followed by a 1:10, 1:100, and 1:1000 dilution of whole blood in sterile PBS. Control diluent (PBS) samples were plated alongside specimens in each case. None of the AKI-or CKD-control mice had evidence of bacteremia. In AKI mice, 1/5 (20%) of the low-dose CS animals were bacteremic. *Among the N=10 AKI mice receiving high dose CS, all 10 suffered mortality (N=3 before collection, among the N=7 remaining, only 5 underwent plating for bacterial culture and all were positive. N=2 of the remaining collected mice were severely ill and blood specimens were prioritized for other biological measurements, thus 5/10 represents 5/5 of the actual plated samples). **Among CKD mice, 2/5 72h-designated animals were euthanized before 72h (one at 36h, one at 48h), and were the identified bacteremic animals.

**Supplemental Figure 2. tGFR changes in sepsis studies.** Absolute tGFR measurements in AKI studies (A) and CKD studies (B). Sham, CS-low dose, and CS-high dose (left, middle, right panels, respectively) studies are shown with baseline and either 24h or 72h followup tGFR. C) Pictured are No-AKI (upper graphs) and AKI (lower graphs) 24-hour animal tGFR curves. Examples of graphs from both sham and high-dose CS animal are shown in comparison. In No-AKI: sham animals, a typical pattern of FITC-sinistrin high amplitude peak after retroorbital injection was appreciated, followed by a decline near background fluorescence skin levels after 1:05h. In the AKI: sham mouse there is a slower decline from peak amplitude and it does not return to background levels by the end of the measurement period. 24-hour follow up tGFR curves are overlayed on the same animal, demonstrating little change in tGFR from baseline to 24 hours in either group receiving sham injections. Conversely, there is significant flattening out of 24-hour follow up tGFR curves in animals receiving high CS doses. In AKI mice, severe renal failure was observed with almost complete plateau of FITC-sinistrin signal, indicating little functional renal clearance. Note the relatively muted signal in AKI-high dose 24-hour curve vs all other curves, attributed to moribund state and little observed animal activity during tGFR measurement period. Within No-CKD group: (# p<0.05 baseline vs 24h No-AKI and No-CKD controls; †p<0.05, ††p<0.01 for baseline vs followup in AKI mice)

**Supplemental Figure 3. Body-weight adjustment CS AKI and CKD Experiments.** (A) Body weight (BW) dose adjustments in AKI groups did not reverse differences in sepsis severity between controls and AKI mice as measured by mortality (B) and clinical sepsis severity (C). In CKD groups, the delivered CS dose was highest in the No-CKD group (D), and two animals suffered severe sepsis and needed euthanasia in this group (E). BW adjustment partially reversed sepsis severity in CKD mice (F), though higher attrition than No-CKD mice was appreciated, again in a delayed fashion. Flat low dose and high dose CS studies are included in AKI and CKD BW data in background for comparison.

**Supplemental Figure 4. Modified Clinical Sepsis Score.** Adopted by Shrum et al.^38^ Animals were graded on a scale of 0 – 4 using individual metrics listed at t=0h (baseline), 4h, 24h, (and 48h, 72h in those designated groups). Total score from all metrics was added for composite score. Rectal temperatures were also recorded alongside sepsis scores.

