## Supplemental Figures for "Title: AKI and CKD Worsen Outcomes in Experimental Sepsis: Creating a Reverse-translational Model of Acute Peritoneal Infection and Recovery in an Immunocompromised Rodent"

**A.**

| | Sham | | Low Dose | | High Dose | |
|---|---|---|---|---|---|---|
| | 24h | 72h | 24h | 72h | 24h | 72h |
| No-AKI | - | - | - | - | - | - |
| AKI | - | - | 1/5 | - | 5/10* | n/a |
| No-CKD | - | - | - | - | - | - |
| CKD | - | - | - | - | - | 2/5** |

**B.**

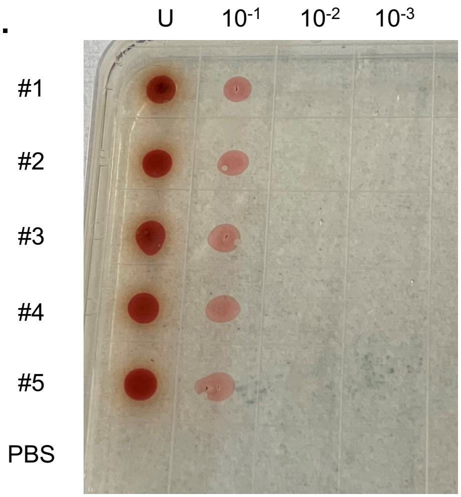


**C.**

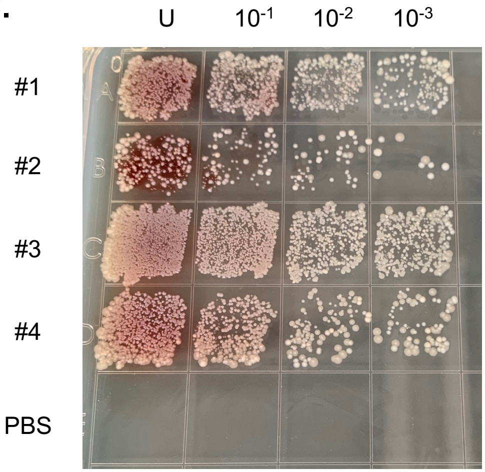

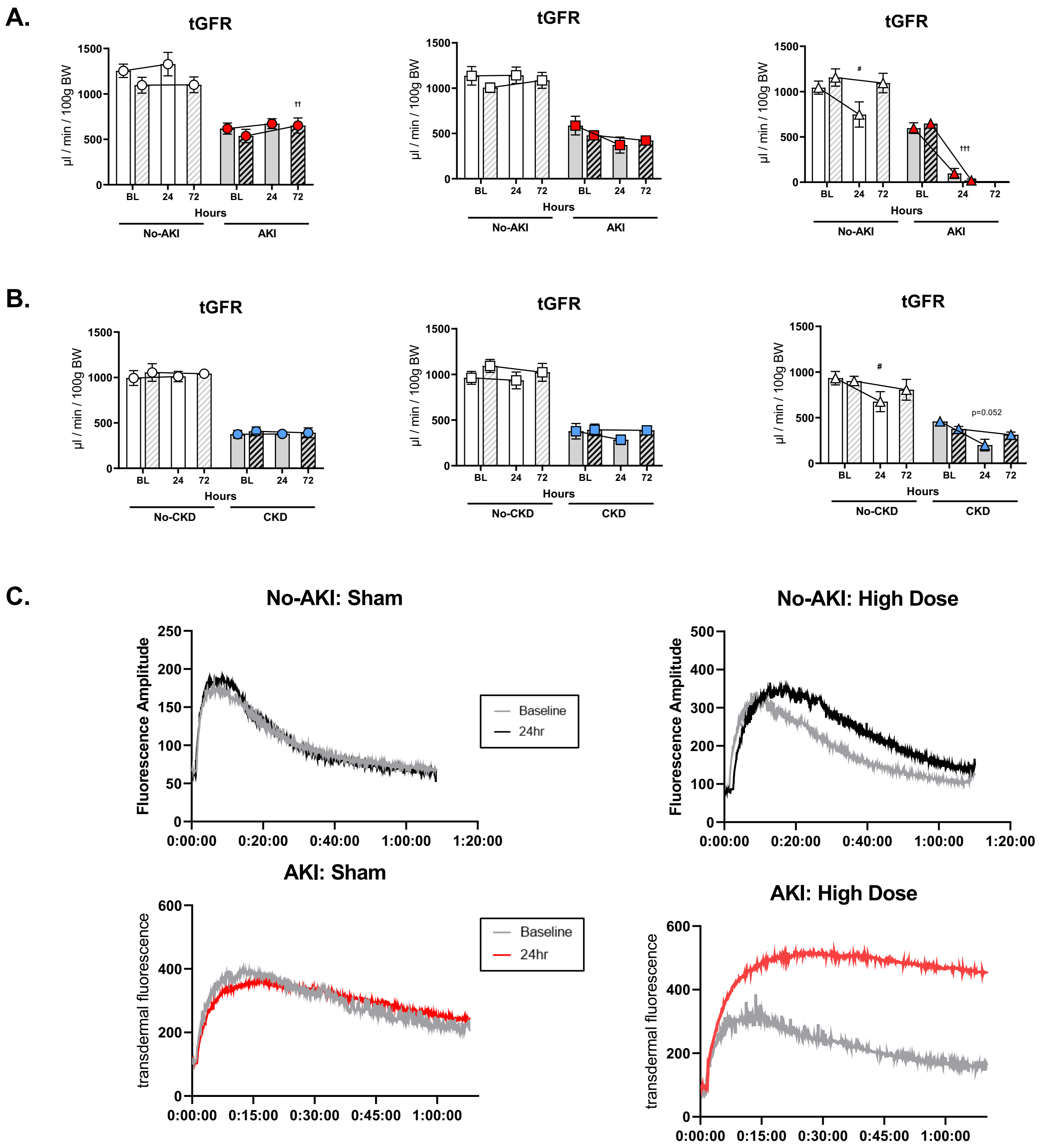

A.
tGFR
µl / min / 100g BW
1500
1000
500
0
BL
24
72
Hours
No-AKI
AKI
††
#
†††
B.
tGFR
No-CKD
CKD
p=0.052
C.
No-AKI: Sham
Fluorescence Amplitude
250
200
150
100
50
0
0:00:00
0:20:00
0:40:00
1:00:00
1:20:00
Baseline
24hr
No-AKI: High Dose
500
400
300
AKI: Sham
transdermal fluorescence
600
0:15:00
0:30:00
0:45:00
AKI: High Dose

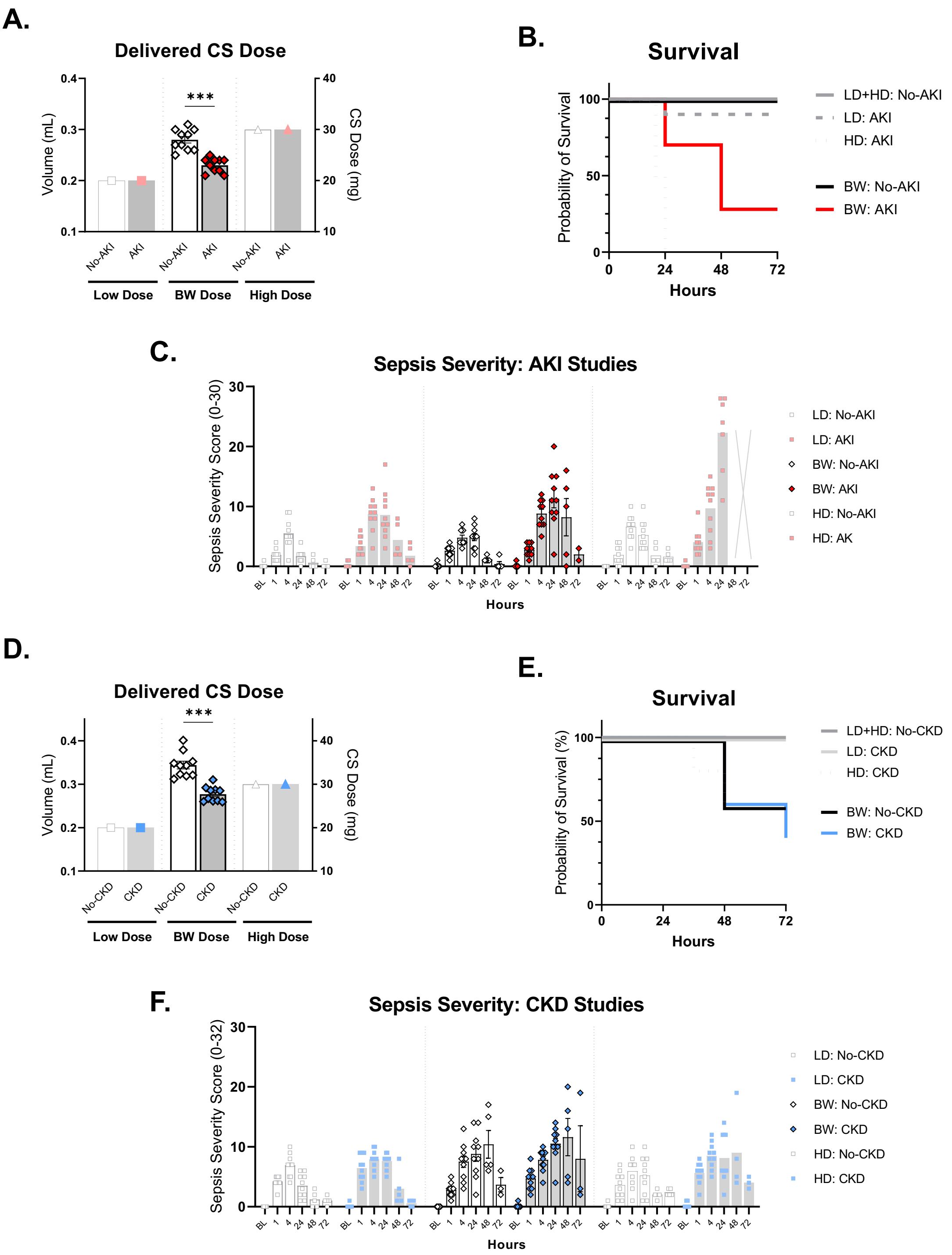
A.
Delivered CS Dose
Volume (mL)
CS Dose (mg)
***
No-AKI
AKI
Low Dose
BW Dose
High Dose
B.
Survival
Probability of Survival
Hours
LD+HD: No-AKI
LD: AKI
HD: AKI
BW: No-AKI
BW: AKI
C.
Sepsis Severity: AKI Studies
Sepsis Severity Score (0-30)
BL
Hours
LD: No-AKI
LD: AKI
BW: No-AKI
BW: AKI
HD: No-AKI
HD: AK
D.
Delivered CS Dose
Volume (mL)
CS Dose (mg)
***
No-CKD
CKD
Low Dose
BW Dose
High Dose
E.
Survival
Probability of Survival (%)
Hours
LD+HD: No-CKD
LD: CKD
HD: CKD
BW: No-CKD
BW: CKD
F.
Sepsis Severity: CKD Studies
Sepsis Severity Score (0-32)
BL
Hours
LD: No-CKD
LD: CKD
BW: No-CKD
BW: CKD
HD: No-CKD
HD: CKD

| Experiment: | | | 1 | 4 | 24 | 48 | 72 | 1 | 4 | 24 | 48 | 72 | 1 | 4 | 24 | 48 | 72 | 1 | 4 | 24 | 48 | 72 | 1 | 4 | 24 | 48 | 72 |
|---|---|---|---|---|---|---|---|---|---|---|---|---|---|---|---|---|---|---|---|---|---|---|---|---|---|---|---|
| Coat | 0 | Smooth coat | | | | | | | | | | | | | | | | | | | | | | | | | |
| | 1 | Patches of hair piloerected | | | | | | | | | | | | | | | | | | | | | | | | | |
| | 2 | Majority of back piloerected | | | | | | | | | | | | | | | | | | | | | | | | | |
| | 3 | "Puffy mouse" | | | | | | | | | | | | | | | | | | | | | | | | | |
| | 4 | Emaciated | | | | | | | | | | | | | | | | | | | | | | | | | |
| LOC | 0 | Alert | | | | | | | | | | | | | | | | | | | | | | | | | |
| | 1 | Alert, avoids upright stance | | | | | | | | | | | | | | | | | | | | | | | | | |
| | 2 | Alertness slow, ambulates | | | | | | | | | | | | | | | | | | | | | | | | | |
| | 3 | Slow, barely moves | | | | | | | | | | | | | | | | | | | | | | | | | |
| | 4 | Stationary | | | | | | | | | | | | | | | | | | | | | | | | | |
| Activity | 0 | Normal activity | | | | | | | | | | | | | | | | | | | | | | | | | |
| | 1 | Slight suppressed activity | | | | | | | | | | | | | | | | | | | | | | | | | |
| | 2 | Sig supressed activity | | | | | | | | | | | | | | | | | | | | | | | | | |
| | 3 | No activity, stationary | | | | | | | | | | | | | | | | | | | | | | | | | |
| | 4 | No activity + tremors | | | | | | | | | | | | | | | | | | | | | | | | | |
| Stimuli | 0 | Resp. to audio/tactile stim. | | | | | | | | | | | | | | | | | | | | | | | | | |
| | 1 | Slow resp. to stimuli | | | | | | | | | | | | | | | | | | | | | | | | | |
| | 2 | No resp. to aud, few touch | | | | | | | | | | | | | | | | | | | | | | | | | |
| | 3 | No resp. to aud, min touch | | | | | | | | | | | | | | | | | | | | | | | | | |
| | 4 | No resp., cannot right self | | | | | | | | | | | | | | | | | | | | | | | | | |
| Eyes | 0 | Open, no secretions | | | | | | | | | | | | | | | | | | | | | | | | | |
| | 1 | Not fuly open +/-min secr. | | | | | | | | | | | | | | | | | | | | | | | | | |
| | 2 | Half closed +/-mod. secr. | | | | | | | | | | | | | | | | | | | | | | | | | |
| | 3 | Nearly closed +/-mod. secr. | | | | | | | | | | | | | | | | | | | | | | | | | |
| | 4 | Closed or milky | | | | | | | | | | | | | | | | | | | | | | | | | |
| Resp | 0 | Normal, rapid resp. | | | | | | | | | | | | | | | | | | | | | | | | | |
| rate | 1 | Slight decreased resp. | | | | | | | | | | | | | | | | | | | | | | | | | |
| | 2 | Moderately reduced resp. | | | | | | | | | | | | | | | | | | | | | | | | | |
| | 3 | Severely reduced resp. | | | | | | | | | | | | | | | | | | | | | | | | | |
| | 4 | Extremely red. resp. | | | | | | | | | | | | | | | | | | | | | | | | | |
| Resp | 0 | Normal | | | | | | | | | | | | | | | | | | | | | | | | | |
| quality | 1 | Brief Labored breathing | | | | | | | | | | | | | | | | | | | | | | | | | |
| | 2 | Labored, no gasping | | | | | | | | | | | | | | | | | | | | | | | | | |
| | 3 | Labored with int. gasps | | | | | | | | | | | | | | | | | | | | | | | | | |
| | 4 | Gasping | | | | | | | | | | | | | | | | | | | | | | | | | |
| | | Time (__0h__) Total Score | | | | | | | | | | | | | | | | | | | | | | | | | |
| | | Time (__1h__) Total Score | | | | | | | | | | | | | | | | | | | | | | | | | |
| | | Time (__4h__) Total Score | | | | | | | | | | | | | | | | | | | | | | | | | |
| | | Time (__24h__) Total Score | | | | | | | | | | | | | | | | | | | | | | | | | |
| | | Time (__48h__) Total Score | | | | | | | | | | | | | | | | | | | | | | | | | |
| | | Time (__72h__) Total Score | | | | | | | | | | | | | | | | | | | | | | | | | |
